# Modulation of cannabinoid receptor signaling by endocannabinoids

**DOI:** 10.1101/2022.08.06.502185

**Authors:** Kaavya Krishna Kumar, Michael J. Robertson, Elina Thadhani, Haoqing Wang, Carl-Mikael Suomivuori, Alexander S. Powers, Lipin Ji, Spyros P. Nikas, Maria Gerasi, Kiran Vemuri, Ron O. Dror, Asuka Inoue, Alexandros Makriyannis, Georgios Skiniotis, Brian Kobilka

**Author notes:** Correspondence (A.M.), (G.S.) and (B.K.K). Equal Contribution (Kaavya Krishna Kumar, Michael J. Robertson).

## Abstract

Endocannabinoids (eCBs) are endogenous lipid molecules that activate the cannabinoid receptor 1 (CB1), a G protein coupled receptor (GPCR) that signals primarily through the G_i/o_ family of G proteins to regulate neurotransmitter release. Consequently, CB1 is an important therapeutic target for several neurological disorders. How eCBs interact with CB1 is not known and the downstream signaling they activate is not well understood. In this study we show that eCBs do not activate G_i_1 as much as synthetic cannabinoids. To characterize activation of CB1 by eCB, we formed an eCB analogue-bound (AMG315) CB1-G_i_ signaling complex for structural studies. The structure reveals differences in the orthosteric ligand binding pocket not seen in the previous CB1 structures, providing insights into the structural determinants of ligand efficacy. In combination with signaling and simulation data, this study provides mechanistic insights into CB1 activation by different classes of ligands, and sheds light on the G protein preferences between endogenous and exogenous ligands.

## Introduction

The cannabinoid receptor type 1 (CB1) is a critical component of the endocannabinoid system and the most abundantly expressed G protein-coupled receptor (GPCR) in the brain^1^. As CB1 regulates a wide range of neuronal functions, it is an attractive target for treating pain, anxiety, anorexia, and neurodegenerative disorders^2–4^. Endogenously, CB1 is activated by two endocannabinoids (eCBs), arachidonoyl ethanolamine (anandamide) and 2-arachidonoyl-*sn*- glycerol (2-AG), that are derivatives of arachidonic acid. CB1 is also activated by many structurally diverse exogenous ligands, most notably, the plant-derived classical cannabinoid (-)-Δ^9^-tetrahydrocannabinol (Δ^9^-THC), the non-classical cannabinoids, exemplified by CP- 55,940, and synthetic cannabinoid receptor agonists (SCRAs) that have emerged as illicit, designer drugs of abuse. Apart from orthosteric agonists, allosteric modulators of CB1 have also been developed recently.

Despite their promising therapeutic potential, exogenous CB1 agonists have a small therapeutic window as they elicit on-target side effects, including catalepsy, hypolocomotion, and memory impairment. Additionally, chronic CB1 activation by orthosteric agonists leads to tolerance and dependence. In addition to these side-effects, SCRA use is associated with more severe side-effects that may even result in death. There is increasing evidence that these severe side-effects caused by SCRAs could be a result of the super-efficacious activation of CB1 signaling that might lead to erratic neurotransmitter modulation and toxicity. On the other hand, positive allosteric modulators (PAMs) of CB1 have shown efficacy in enhancing the antinociceptive effects of endocannabinoids *in vivo* without any cardinal signs of CB1 side-effects or tolerance^5^.

At the cellular level, CB1 predominantly signals through the adenylate cyclase inhibitory G protein family, G_i/o_, and also recruits arrestins. Ligands are pleiotropically coupled to multiple signaling pathways and can stabilize conformations that might favor interaction with specific effectors. As with other GPCRs, ligands biasing the receptor towards interactions with specific G_i/o_ subtypes or arrestins may exhibit differential behavioural outcomes, thus increasing the therapeutic window. Therefore, a better understanding of the structural basis of CB1 activation with diverse ligands could offer valuable insight and enhance our ability to design novel drugs with improved pharmacological profiles.

We have previously determined the structure of CB1 bound to a SCRA, FUB^6^ and others have determined CB1 structures bound to the classical cannabinoid analogues AM841 and AM11542^7^, the non-classical cannabinoid CP-55,940 and the negative allosteric modulator (NAM) Org27569^8^. However, no structure of CB1 bound to an eCB is available. To understand the structural basis of CB1 activation by eCBs, we determined a 3.4 Å cryo-EM structure of an eCB analogue-activated full length CB1 in complex with heterotrimeric G_i1_ protein. We employed a metabolically stable and potent anandamide analogue, AMG315, for structure determination. To gain a better understanding of the downstream pathways activated by the ligands (eCBs, phytocannabinoids and synthetic cannabinoids), we used fluorescence spectroscopy and signaling assays to show that different cannabinoids activate G_i_1 to different extents. Further, with molecular dynamics (MD) simulations, mutagenesis and signaling data we glean insights into the structural determinants of ligand efficacy in CB1.

## Results and Discussion

### Activation of G_i_1

CB1 preferentially signals via the G_i/o_ G protein subtype. To investigate how structurally diverse ligands activate G_i_1, we performed a GTP turnover assay using the non-classical CP55,940 (CP) and the synthetic cannabinoid MDMB-Fubinaca (FUB), the eCB anandamide, as well as AMG315, an eCB analogue that has two carefully chosen chiral centers and exhibits remarkable biological activity and stability^9^ (Fig 1a). The full agonists CP and FUB were equally efficacious towards G_i_1 (Fig 1b). When compared to CP and FUB, AMG315 was slightly less efficacious for G_i_1 (Fig 1b). On the other hand, anandamide acted as a weak partial agonist for G_i_1 (Fig 1b) and was able to induce only 60 % GTP-turnover, compared to CP and FUB in G_i_1.

**Fig 1.**
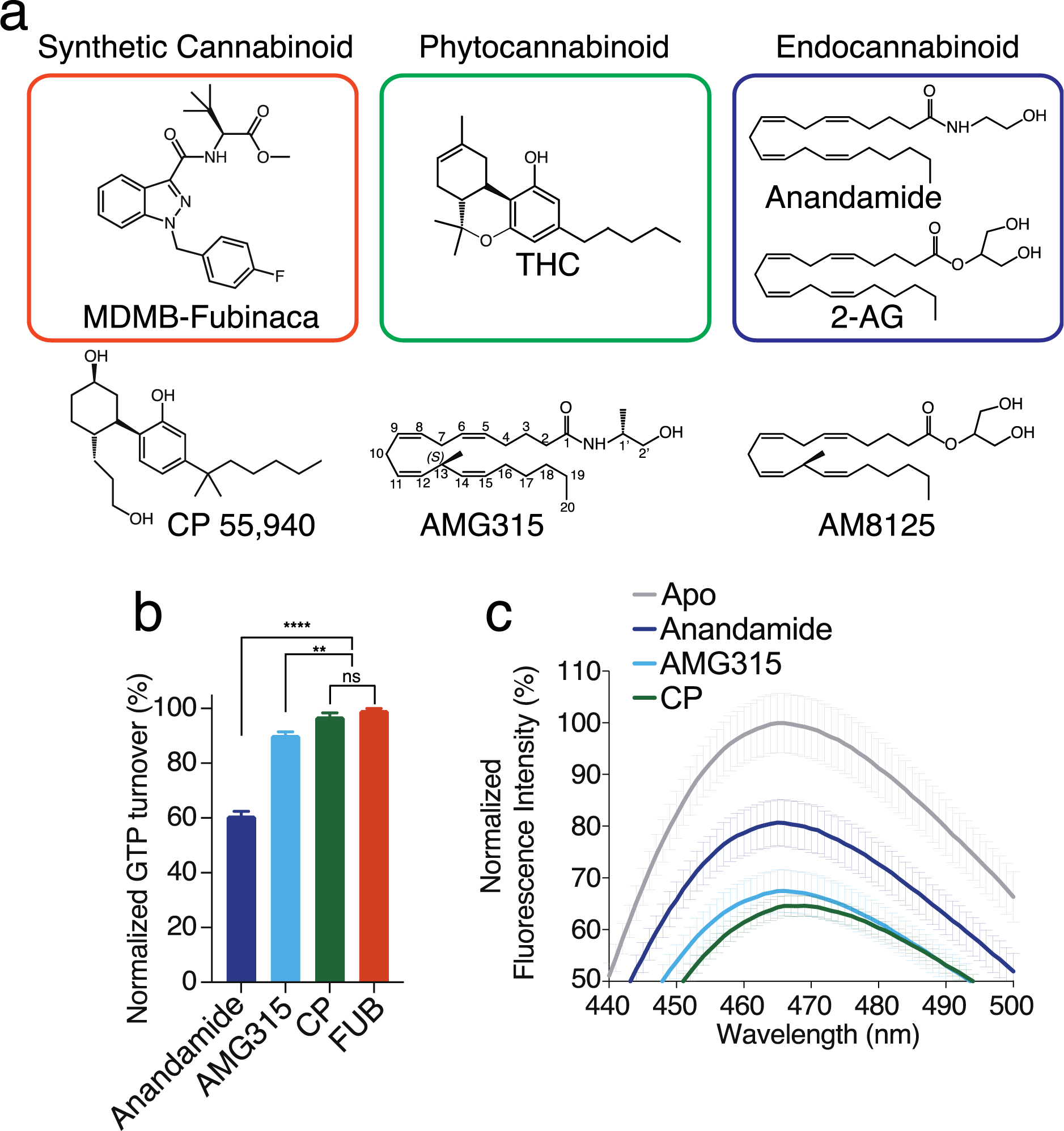
a. Chemical Structures of a synthetic cannabinoid (FUB), Phytocannabinoid (THC) and endocannabinoids (Anandamide and 2-AG). Structures of analogues of phytocannabinoid (CP 55, 940) and endocannabinoid (AMG315 and AM8125) used in this study. b. GTP turnover assay showing efficient turnover produced by CP, AMG315 and anandamide with G_i1_ (Data normalization done with FUB as 100% and receptor alone as 0%). (Mean ± SD, *p* < 0.0001**** and *p <* 0.01**, *t*-test) c. Bimane spectra monitoring TM6 showing differences between anandamide-bound CB1 compared to with CP and AMG315.

To understand how ligands of different efficacies stabilize TM6, we performed fluorescence spectroscopy with CB1 labeled with the environmental-sensitive fluorophore, monobromobimane (bimane) as a conformational reporter of TM6 activation of CB1. To enable site-specific labelling, a minimal cysteine version of CB1 was generated^10^ where all the cysteine residues (except C256 and C264 that form disulfide) were mutated to alanine. A cysteine residue was engineered at residue 241 (L6.33) on TM6, which was labelled with bimane. The bimane spectra of CP- and AMG315-bound CB1 were not significantly different except for a small (1nm) blue-shift in λ_max_ (Fig 1c). Adding Anandamide to CB1, on the other hand, results in a smaller decrease in intensity and a blue-shift in λ_max_ by 4 nm compared to CP (and 3 nm compared to AMG315) (Fig 1c). These differences in the bimane spectra perhaps indicates that anandamide stabilizes a distinct conformation in TM6.

### Determination of an endocannabinoid-bound CB1-G_i_1 complex

To better understand the structural differences in the ligand binding mode of eCBs compared to SCRAs like FUB^6^, and the classical cannabinoid AM841^7^, we determined the structure of an eCB-bound CB1 signaling complex. In order to determine the structure of CB1 bound to a ligand with an eCB-like chemical structure, we tested eCB (both anandamide and 2-AG) analogues for their ability to induce CB1-dependant G_i_1 GTP turnover as a measure of complex formation and stability. The anandamide analogue AMG315 and the 2-AG analogue AM8125 (Fig. 1a) induced significantly better GTP turnover compared to their parent compounds (Fig. 2a). However, as expected, neither (AMG315 and AM8125) were more efficacious than FUB or the THC analogue, CP (Fig. 2a). Size exclusion chromatography (SEC) showed that CB1 formed a slightly more stable complex with AMG315 than AM8125 (Fig. S1a). The PAM ZCZ-011 (ZCZ) was able to further improve GTP turnover and further stabilize the complex (Fig. 2b). CB1 bound to AMG315 and ZCZ formed a complex with G_i_1 that was stable enough for cryoEM imaging yielding a density map at a global nominal resolution of 3.4 Å (Fig. S1b). In order to test if we could observe better density for a CB1 PAM, we also determined the structure of AMG315-bound CB1-G_i_1 complex in the presence of AM11517, that showed better PAM activity (Fig S1c) and yielded a 3.1 Å resolution map (Fig. S1d). We observed slightly different poses for AMG315 in the two maps and as a result some differences in the ligand binding pocket (discussed later). Since, otherwise the maps are very similar, we use only the ZCZ-bound map (as our previous FUB-bound structure was also obtained in the presence of ZCZ) for further discussions. Recently, structures of CB1 bound to ZCZ were determined, showing binding site involving TMs 2, 3 and 4^11^. In neither of our structures do we see any density in the region described in the previous study^11^ and hence, we do not model either of the PAMs (ZCZ and AM11517) in our structures. Overall, the mode of G_i_ engagement with the endo-bound CB1 is very similar to the previously determined structure of FUB-bound CB1 complex, except for ∼ 4 Å deviation of the αN of G_i_ between the CB1-G_i_ complexes (Fig S2a).

**Fig 2.**
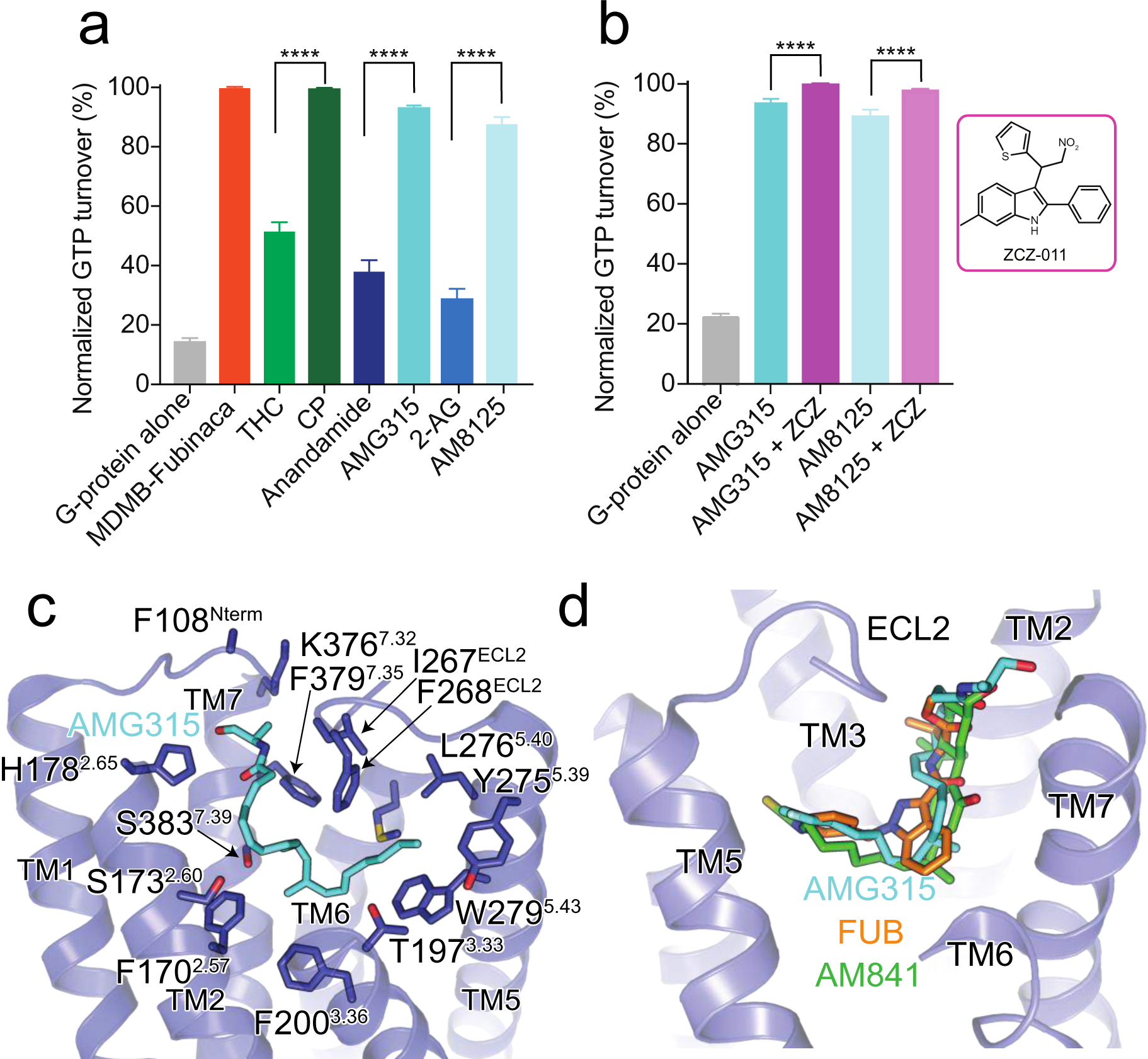
a. GTP turnover assay with G_i_1 showing maximum turnover produced by FUB and CP. Endocannabinoid produce much lower GTP turnover compared to their analogues. (Mean ± SD) b. Addition of the PAM, ZCZ increases GTP turnover of the endocannabinoid analogues. (Mean ± SD, *p* < 0.0001****, *t*-test) c. AMG315 binding pocket showing residues that are within 4 Å from the ligand. d. Overlay of ligands from different chemical classes, synthetic cannabinoid (FUB, orange), classical cannabinoid (AM841, green) and endocannabinoid (AMG315, blue).

### Endocannabinoid Interaction

The cryoEM map shows well-defined density that allows unambiguous modelling of AMG315 and all the protein components of the CB1-G_i_1 complex (Fig. S1b, d, Fig. S2b). AMG315 engages the receptor through hydrophobic and polar interactions (Fig. 2c). The acyl chain of AMG315 is buried deep in the binding pocket, while the polar head group is closer to the extracellular pocket, interacting with the ‘lid-like’ N-terminus (Fig. 2c) and pointing into a largely positive cavity formed by the TM1-TM7 interface (Fig. S2c). Previously we observed a phospholipid molecule bound in the TM1-TM7 interface during MD simulations^6^. Phospholipids are the precursors for endocannabinoids and lipid binding observed in the simulations might indicate the ligand entry point for endocannabinoid through the membrane. The ligand access point in the TM1-TM7 interface is positively charged while the rest of the binding pocket is largely uncharged (Fig. S2c). This charge distribution might help align the acyl chain and the hydroxyl of the endocannabinoid head group and guide the ligand correctly into the binding pocket. This mechanism of guided ligand entry has been proposed for other lipid binding GPCRs such as S1P1 and LPA1^12^.

AMG315 overlays well with the previously determined structures of distinct classes of cannabinoids such as FUB^6^ and AM841^13^ (Fig. 2d). In addition to most of the hydrophobic and polar interactions made by FUB and AM841, AMG315, through its carbonyl head group, interacts with N-terminal residue F108^Nterm^ and I267^ECL2^ in ECL2 (Fig. 2c). When comparing CB1 and CB2, the N-terminus and ECL2 are the most diverse regions in terms of length and sequence conservation. The interactions of AMG315 with these regions might explain its unprecedented 20-fold selectivity for CB1 over CB2^9^.

The residues, F200^3.36^ and W356^6.48^ (known as the “toggle switch”), play an important role in stabilizing the inactive conformation of CB1^14^, wherein F200^3.36^ and W356^6.48^ form π-π aromatic stacking interactions (grey, Fig. 3a). Since these two residues are important for CB1 signaling activity, we speculated that a ligand’s efficacy correlates with its ability to engage the “toggle switch” to activate CB1. Upon activation, the rotation of TM3 and TM6 disrupts the stacking of F200^3.36^ and W356^6.48^ (Fig. 3a), with the phenyl ring of F200^3.36^ pointing towards the ligand to form hydrophobic interactions. In the FUB-bound structure, F200^3.36^ interacts with the indazole ring (Fig. 3b). In the case of AMG315, the methyl group at C-13 (S-stereochemistry) interacts with the “toggle switch” residues (Fig. 3b, black circle).

**Fig 3.**
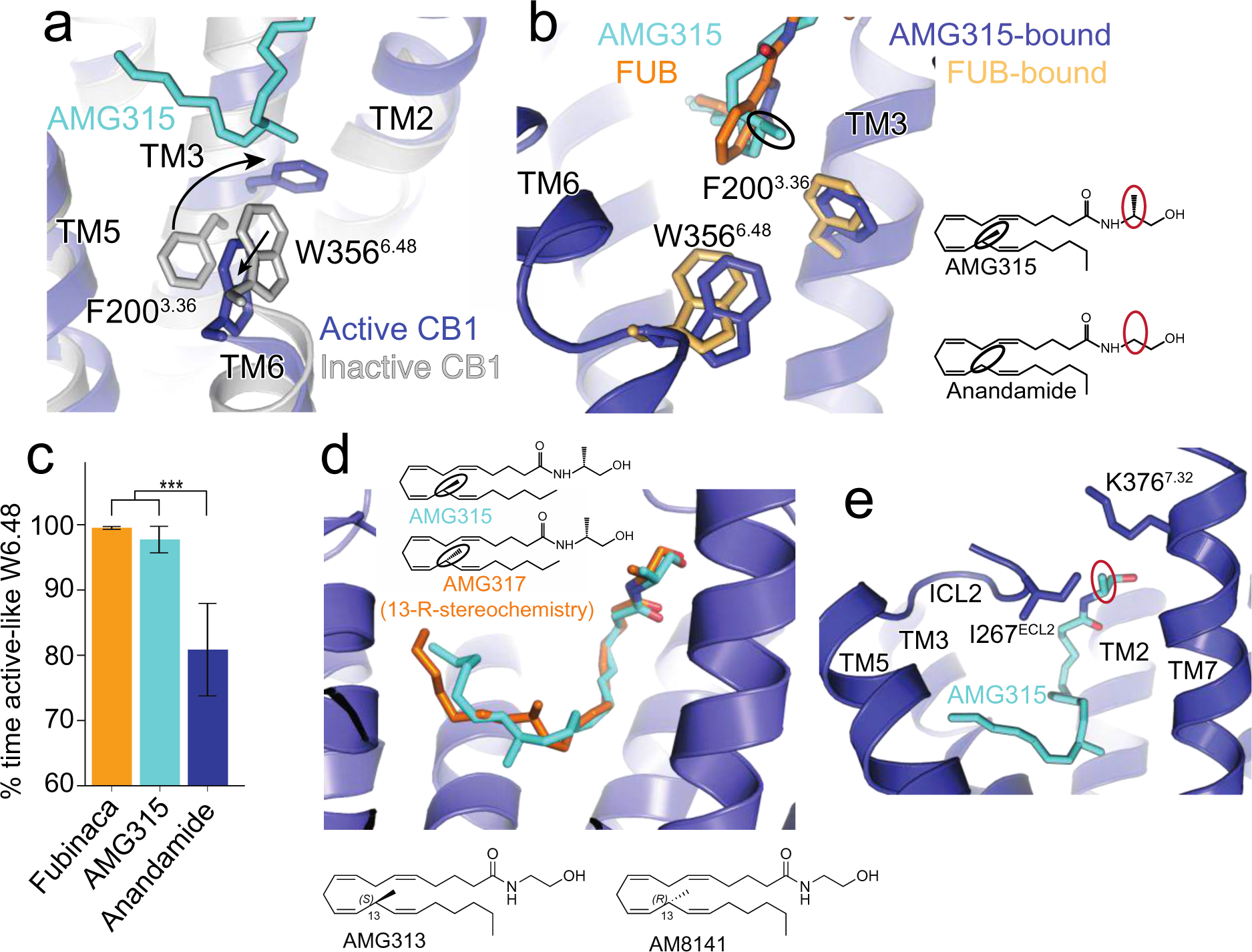
a. AMG315 stabilizes the ‘toggle switch’ residues W356^6.48^ and F200^3.36^ in the active state. b. Ligand interaction with residues W356^6.48^ and F200^3.36^. The methyl group in AMG315 (circled black) not present in anandamide makes interaction with the ‘toggle switch’. c. In simulation, Anandamide stabilizes active-like conformations of W356^6.48^ significantly less when compared individually to FUB (*p* = 0.02, two-sided Welch’s *t*-test) and to AMG315 (p = 0.04), as well as when compared to both FUB and AMG315 as a group (p = 0.003). d. Molecular docking showing an R-enantiomer instead of the S-enantiomer at position 13 in AMG315, repositions the methyl group away from the “toggle switch”. Insert below: chemical structure of position 13 substituents of anandamide, AMG315 (S-enantiomer) and AM8141 (R-enantiomer). e. The methyl group at position 1’ (circled red in 3B) on AMG315 interacts with residues on ECL2 (Ile267^ECL2^) and extracellular region of TM7 (K376^7.32^). This substitution is not found in anandamide.

Molecular dynamic (MD) simulations show that anandamide stabilizes W356^6.48^ in the active-like conformation significantly less than do FUB and AMG315 (Fig. 3c), implying that lower efficacy of anandamide might be related to its ability to interact with and activate the “toggle switch” residues. Molecular docking shows that using AMG317 with the R-stereochemistry at C-13 instead of S-stereochemistry as seen in AMG315, makes the methyl substitution point away from the “toggle switch” (Fig. 3d), and consequently no detectable receptor activation is observed for AMG317 (R-stereochemistry, Fig. 3d)^9^. Furthermore, by comparing anandamide with its (S)-C-13 methyl congener (AMG313, Fig. 3d), we observe that this methyl group, imparts a 5-fold increase in potency and an increase in efficacy. However, the (R)-C-13 methyl enantiomer, AM8141 (Fig. 3d), shows no detectable activity at CB1 ^9^.The other methyl substitution on AMG315 compared to anandamide at C-1’ interacts with residues on ECL2 (Ile267^ECL2^) and the extracellular region of TM7 (K376^7.32^) (Fig. 3e, red circle). The combined interactions of the receptor with AMG315 due to the two chiral methyl groups synergize to result in an increase in potency of over 100-fold compared to anandamide^9^.

### Role of TM2 in ligand efficacy

Agonist interactions with TM2 appear to play a more important role for CB1 activation. As with previous agonist-bound structures, the AMG315-bound CB1 shows extensive structural rearrangements in the ligand binding pocket compared to the antagonist-bound structure^6,13, 15^. Upon AMG315 binding, the N-terminus of CB1 is displaced from the transmembrane core, followed by the inward displacement of TM1 and TM2 (Fig. 4a). This inward movement of TM2 is accompanied by the repositioning of residues F170^2.57^, F174^2.61^, F177^2.64^ and H178^2.65^, that rotate towards and interact with the agonist (Fig. 4b). These structural differences in the ligand binding pocket between binding of agonist and antagonist are not seen in the closely related CB2 receptor (Fig. S3a). Since TM2 rearrangement is stabilized by agonist binding, these differences might be an important determinant of ligand efficacy in CB1. In the previously determined FUB-bound CB1-G_i_1 structure, the *tert*-butyl group of FUB interacts with these repositioned residues on TM2 (Fig. 4c). MMB-Fubinaca, which has an isopropyl substitution at this position, has a reduced efficacy (Fig. S3b) and potency^16^ compared to the *tert*-butyl substituent of FUB indicating that, in addition to interaction with the “toggle switch” residues, TM2-ligand interactions is an important determinant of ligand efficacy. Though AMG315 overlays well with FUB in the ligand binding pocket and makes similar interactions with the receptor, AMG315 is a less efficacious ligand compared to FUB. This difference in efficacy might be attributed in part to the interactions the ligands make with TM2, wherein FUB has more extensive interactions with TM2 than does AMG315 (residues that are further than 4 Å for the AMG315-bound structure are shown as light blue, Fig. 4c). The residues on TM2 that are within 4 Å of FUB are F170^2.57^, S173^2.60^, F174^2.61^, F177^2.64^ and H178^2.65^ (Fig. 4c). However, only residues F170^2.57^, S173^2.60^ and H178^2.65^ are within 4 Å of AMG315 (Fig. 4c). Studies have shown that adding a methyl substitution in anandamide at C-7 (AM11604, Fig. S3c) increases the efficacy (E_max_) to 100% relative to the full agonist CP55940^9^, presumably due to its enhanced interactions with residues of TM2. Compared to anandamide (E_max_ 61%), AMG315 with a methyl substitution at C-13 has an E_max_ value of 76% probably due to the gained interaction with the “toggle switch” residues (discussed earlier). However, introducing a substitutions at C-7 (known as AM11605, Fig. S3c), which would gain interactions with TM2, increases the E_max_ value to 100%^9^.

**Fig 4.**
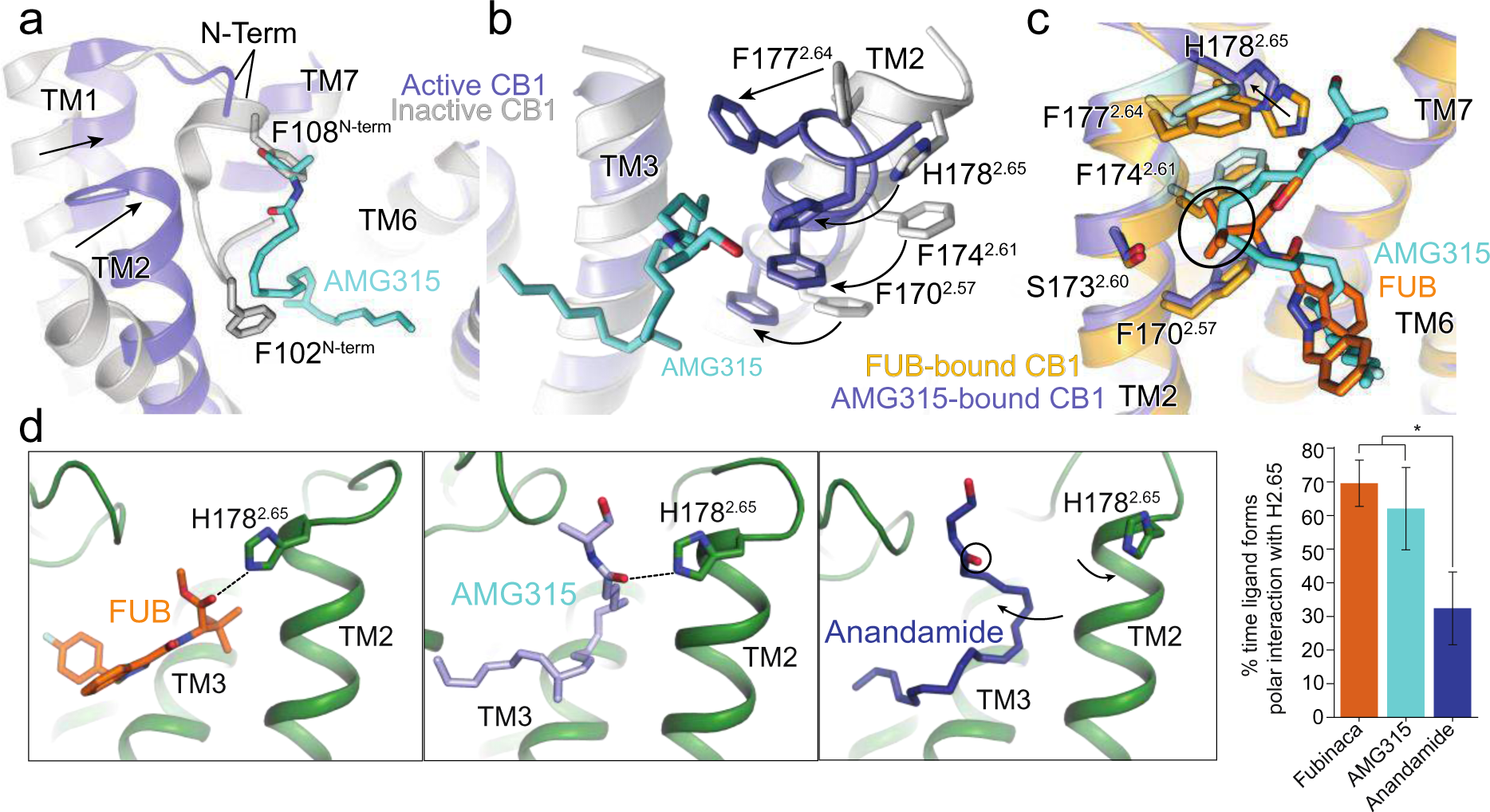
a. Overlay of inactive structure of CB1 (PDB: 5U09, white) and AMG315-bound structure (blue) showing the inward movement of TM2 upon activation and displacement of the N- terminus. b. The inward movement of TM2 from the inactive state (grey) to the active state (blue) results in the translocation of residues F177^2.64^, H178^2.65^, F174^2.61^ and F170^2.57^ towards the agonist. c. FUB interacts with more TM2 residues (orange) compared to AMG315 (blue). F177^2.64^ and F174^2.61^ interact with FUB but not AMG315, shown in light blue. d. In simulation, FUB and AMG315 form polar interactions with H178^2.65^ more often than the less efficacious partial agonist Anandamide (*p* = 0.01, two-sided Welch’s *t*-test).

Compared to the FUB-bound structure, in the AMG315-bound CB1, H178^2.65^ has moved away from the ligand by ∼ 2.5 Å (Fig. 4c). To investigate if there is a correlation between ligand efficacy and interaction with H178^2.65^, we performed MD simulations to probe the frequency of interactions between ligands (FUB, AMG315 and anandamide) and H178^2.65^. The full agonist FUB and the slightly less efficacious AMG315 more often form a polar interaction with H178^2.65^ compared to the partial agonist anandamide (Fig 4d). As described above, the methyl substitution at C-1’ in AMG315 interacts with K376^7.32^ and Ile267^ECL2^ (Fig. S2e) which might limit its movement in the ligand binding pocket, increasing interaction frequency with H178^2.65^ (Fig. 4d). The absence of this methyl substitution in anandamide would allow the ligand to move away from TM2 and more often break interaction with H178^2.65^. Though not statistically significant, the frequency of polar interaction between H178^2.65^ and AMG315 is lower than with FUB (Fig 4d). In our other structure of the CB1-G_i_ complex (with AM11517 as the PAM), AMG315 is bound in a conformation wherein the carbonyl group of AMG315 has moved away from the H178^2.65^ such that the His residue is not within hydrogen bonding distance (Fig. S3d, Fig. 4d).

### Distinct role of TM2 in activation of CB1

Changes in the extracellular end of TM2 upon agonist binding are associated with changes in the intracellular end of TM2, wherein a group of residues undergo rearrangement upon activation. At the intracellular end of TM2, F155^2.42^ undergoes a concerted movement with F237^4.46^ upon activation. In the inactive structure, the aromatic ring of F237^4.46^ is facing inward, towards TM2-3, with F155^2.42^ positioned at the core of the receptor (Fig. 5a). Upon activation, F237^4.46^ and F155^2.42^ rotate outward away from the receptor core (Fig. 5a). Along with the F155^2.42^, the intracellular side of TM2 rotates with the sidechain of H154^2.41^ moving outward ∼ 4Å compared to the inactive CB1 structure (Fig. 5a). In both the active^17^ and inactive^18^ structures of CB2, F72^2.42^ is positioned outward and is not facing the core of the protein. Hence, the intracellular end of TM2 in CB2 does not undergo a similar rotation as in CB1, and there is little difference in the orientation of positions 2.42 or 4.46 between inactive and active states (Fig. 5b). In fact, this structural change at position 2.42 is not seen in any other receptors including β_2_AR, μOR and M2R (Fig. S4a). This is probably because none of these receptors has a bulky aromatic residue at position 4.46, that would sterically clash with Phe at position 2.42 upon activation (Fig. S4a-b), which might make this mechanism of activation unique to CB1. Mutating F237^4.46^ to a Leu (like in CB2) increased basal activity^19^. This could be due to the inability of a Leu residue at this position to stabilize an inward rotation of F155^2.42^. A caveat in interpreting the position of residues F237^4.46^ and F155^2.42^ from the crystal structure of inactive CB1 is that the structure was determined with an inactivating TM3 mutation T210^3.46^A ^15^ and the conformation of F155^2.42^ (i.e. inward facing) might be influenced by the presence of this mutation (Fig. S4c). Regardless, the inactive structure of CB2 was also determined with this mutation (T^3.46^A) and the reorientation of F72^2.42^ is not seen in this structure.

**Fig 5.**
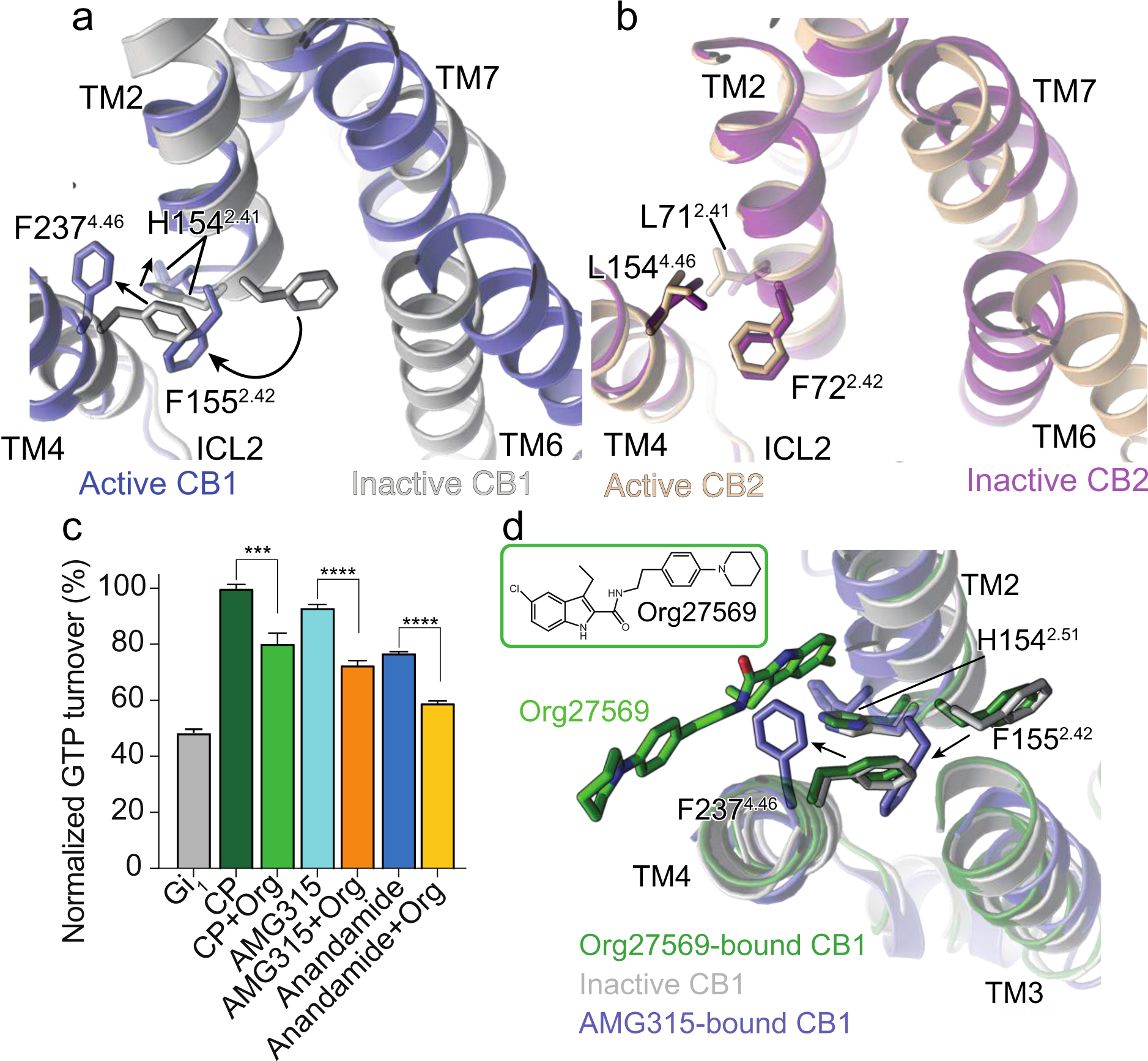
a. Concerted movement of F237^4.46^ and F155^2.24^ upon activation of CB1 from inactive (grey) to active (blue) state. b. The residue at position 4.46 in CB2 is a Leu and does not undergo movement upon activation. c. GTP turnover assay showing reduced turnover in the presence of Org27569 with G_i_1. (Mean ± SD, *p* < 0.0001**** and *p <* 0.001***, *t*-test) d. Structural rearrangement in F237^4.46^ and F155^2.42^ in Org27569 (PDB: 6KQI, green) bound structure compared to active AMG315-bound (blue) and inactive (grey) structures.

CB1 negative allosteric modulator (NAM) Org27569 (Org) has been shown to modulate the F155^2.42^-F237^4.46^ “activation switch” of CB1 to exert its NAM activity. Org decreases GTP turnover in the presence of CP, AMG315 and anandamide (Fig 5c). Org has an unusual pharmacology in that, unlike conventional NAMs, it increases agonist affinity while decreasing G_i_1 turnover. This atypical pharmacology of Org is probably related to the CB1 residues the ligand interacts with. The structure of Org-bound CB1 shows that Org interacts directly with H154^2.41^ in TM2 and stabilizes the inward position of F237^4.46^ (Fig 5d, S5a)^8^, Mutations to F237^4.46^ has been shown to change orthosteric ligand binding. F237^4.46^L mutation increases orthosteric agonist potency^20^ and increases receptor internalization^19^. Mutation of F237^4.46^ to a smaller amino acid Leu, would not cause a steric clash with F155^2.42^, allowing F155^2.42^ to remain in the outward “active” conformation, as mentioned previously. In the WT receptor, Org binding drives the conformational change in F237^4.46^, which somehow causes changes in agonist potency. Change in F237^4.46^ affects the conformation of TM2 and as mentioned previously, the extracellular region of TM2 undergoes a large inward rotation when bound to agonists. Therefore, an explanation for how Org affects agonist potency could be that Org binding increases the propensity of extracellular TM2 to rotate inward, thereby stabilizing an agonist-binding conformation, thus increasing agonist potency, while stabilizing a TM6 conformation that is not conducive for G_i_ binding. For its NAM activity, Org binding stabilizes the inward rotation of F237^4.46^ which probably results in the inward rotation of F155^2.42^ to the receptor core, stabilizing an inactive conformation. In spite of acting as allosteric modulators at CB1, Org shows no activity at the closely related CB2 (Fig S5b)^5, 21, 22^. Org interacts with H154^2.41^ and F237^4.46^ of CB1 and in CB2 the positions 2.41 and 4.46 are both Leu residues (Fig S4a). More importantly CB2 does not contain the F155^2.42^-F237^4.46^ “activation switch” of CB1, and hence, Org is unable to modulate the “activation switch” residues in CB2 to produce their allosteric activity.

## Conclusion

eCB signaling plays a critical role in maintaining homeostasis and is involved in the regulation of neurotransmission and synaptic plasticity. Phytocannabinoids and synthetic cannabinoids that emulate eCB signaling through the CB1 receptor produce undesirable side-effects. Structurally and pharmacologically, eCBs are very distinct from phytocannabinoids and synthetic cannabinoids and understanding signaling by eCBs have important implications for designing drugs with desired signaling profiles.

Anandamide has a lower efficacy compared to the agonist, CP and we show that anandamide might stabilize a distinct conformation of TM6. To better understand the pharmacology of eCBs, we determined the cryoEM structure of a CB1-G_i_ signaling complex bound to AMG315, a metabolically stable and highly potent endocannabinoid analogue. This compound interacts with the N-terminal, TM1 and TM7 regions of CB1 which are not explored by other ligands. Using MD simulations and SAR data, we show that the efficacy of CB1 ligands depends on their propensity to interact with the ‘toggle switch’ residues F200^3.36^/W356^6.48^. Additionally, ligand efficacy in CB1 appears to be related to its interaction with the extracellular end of TM2. Ligand interactions in the extracellular region is transmitted to TM2 intracellular end where residue F155^2.42^ undergoes concerted movement with F237^4.46^ to contribute to activation of CB1. This activation mechanism appears unique to CB1 (not seen in other GPCRs thus far) due to the distinctively positioned Phe residue at position 4.46, and CB1 allosteric modulators appear to regulate this “activation switch” residues to exert their signaling effects.

## Supplementary Figs

**Fig S1.**
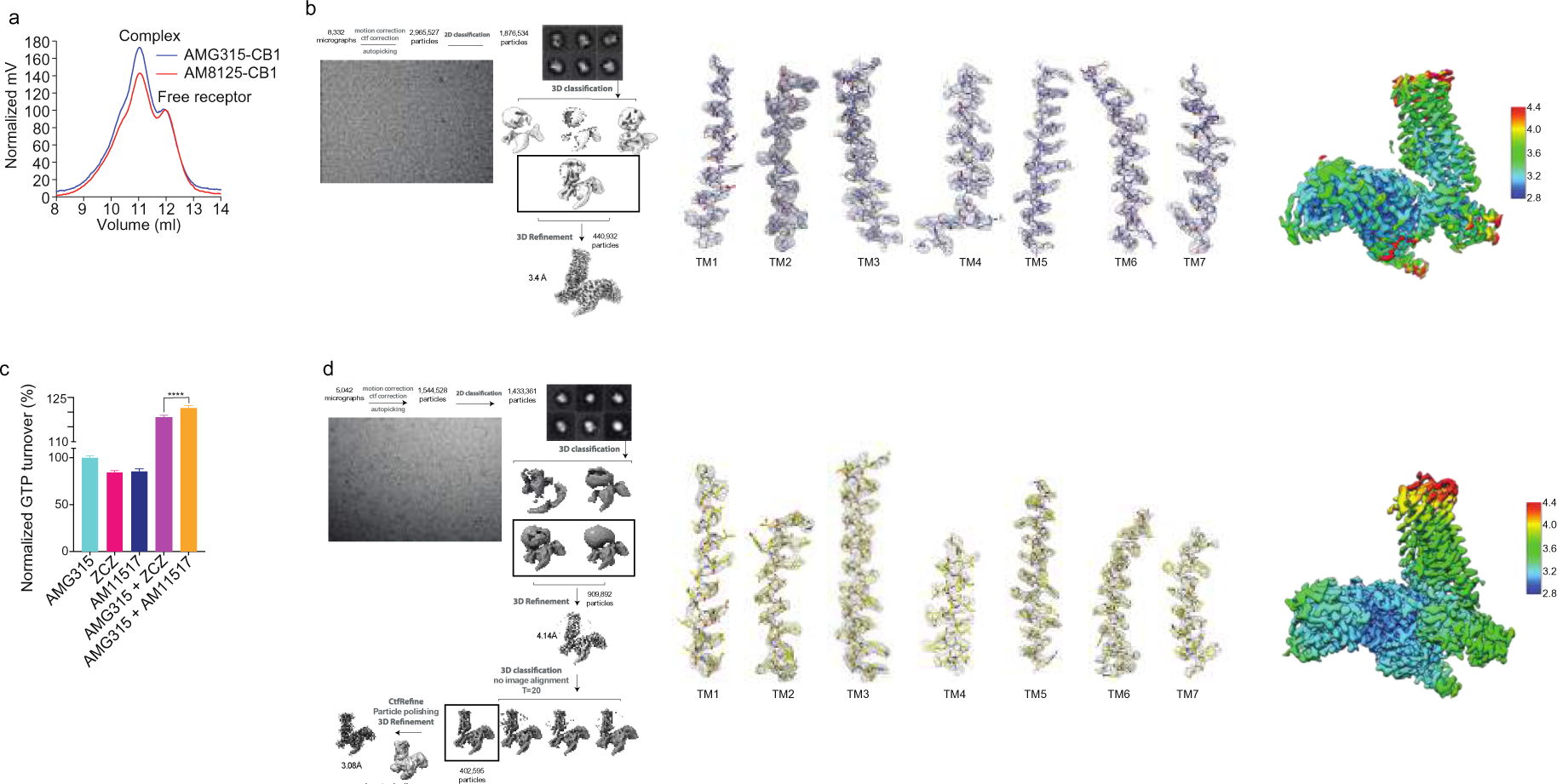
a. Fluorescence-detection size exclusion chromatography (FSEC) traces showing complex peak and free receptor peak with the different AMG315 (blue) and AM8125 (red). b. Cryo-EM processing and map density of TMs in ZCZ-bound CB1 structure. c. GTP turnover assay showing increased turnover with AM11517 compared to ZCZ. d. Cryo-EM processing and map density of TMs AM11517-bound CB1 structure.

**Fig S2.**
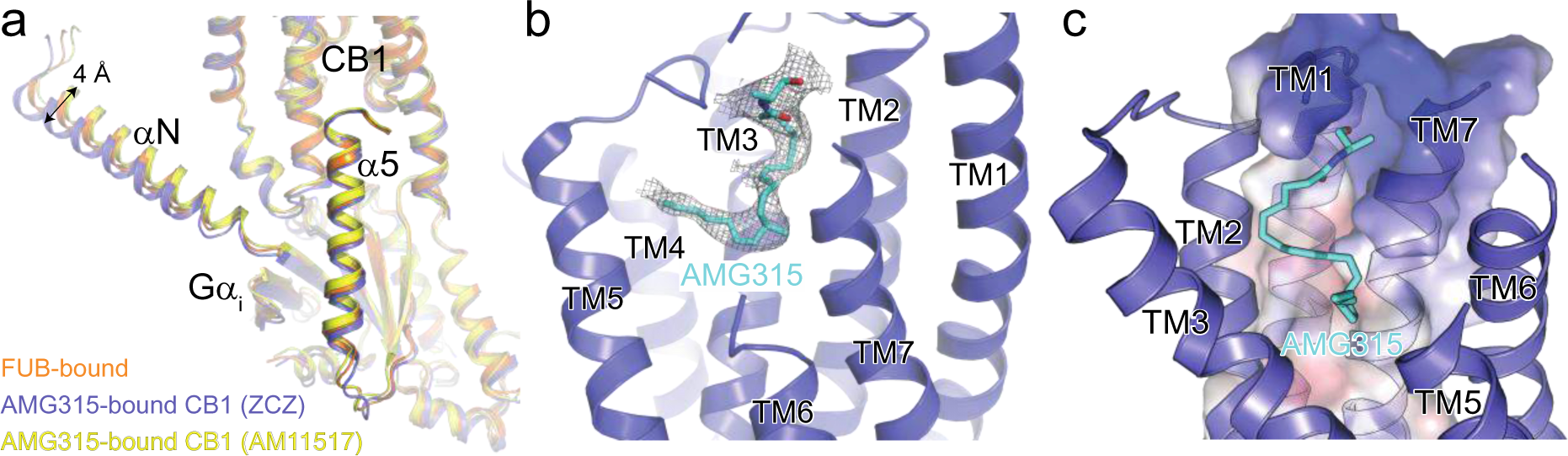
a. Overlay of the Gai subunit from FUB-bound (orange) and the two AMG315-bound (ZCZ, blue and AM11517, yellow), showing a 4Å difference in αN. b. Cryo-EM map density of the orthosteric ligand, AMG315. c. Surface electrostatics (calculated and analyzed by the APBS Electrostatic *PyMol* Plugin with negative colored blue and positive colored red) of the ligand binding pocket formed by TM1 and TM7.

**Fig S3.**
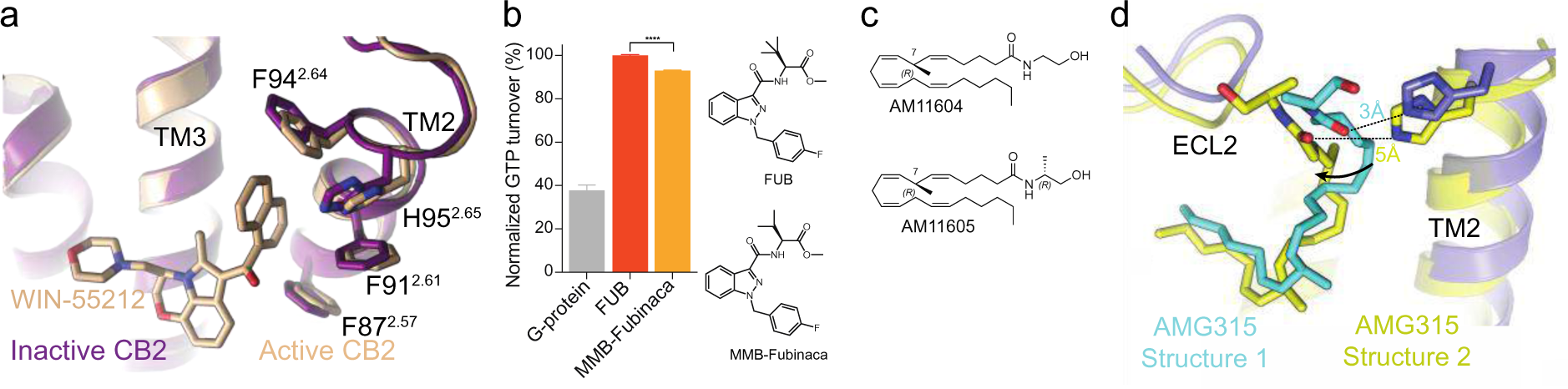
a. Active (PDB: 6PT0) and inactive (PDB: 6KPC) structures of CB2 showing no change in TM2 position upon activation. b. GTP turnover assay showing more turnover with FUB compared to MMB-Fubinaca. (Mean ± SD, *p* < 0.001****, *t*-test) c. Chemical structure of AM11604 and AM11605. d. The difference in conformation in AMG315 between the AMG315-bound CB1 structures, wherein contacts the ligand contact H178^2.65^ in one structure and not the other.

**Fig S4.**
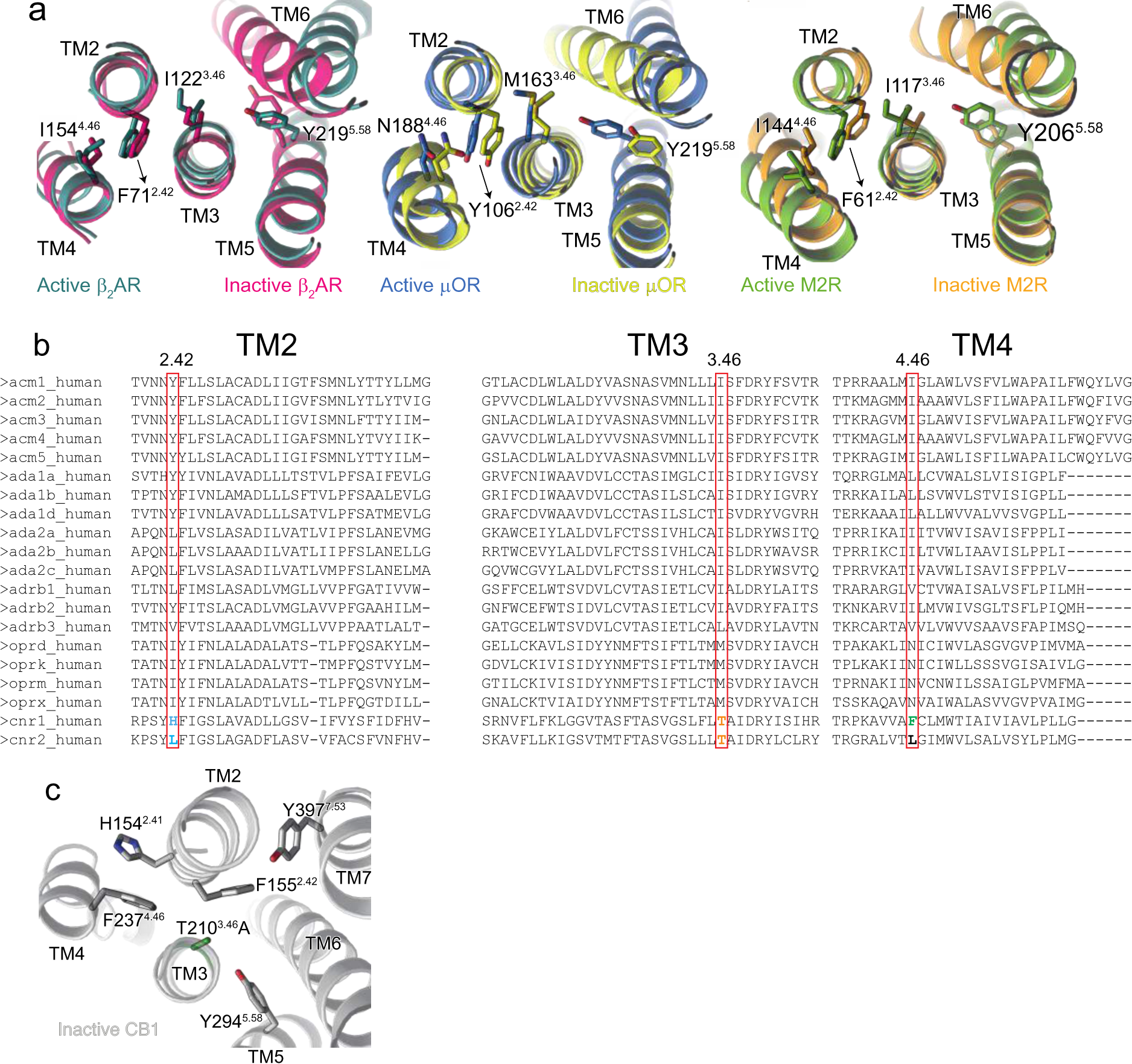
a. Structural changes to TM2-3-4 upon activation in β2AR (Active, PDB code: 3SN6, teal and Inactive PDB: 2RH1, magenta), μOR (Active, PDB code: 6DDE, blue and Inactive PDB: 4DKL, yellow) and M2R (Active, PDB code: 6OIK, green and Inactive PDB: 3UON, orange). b. Alignment of GPCRs showing differences in residues at position 2.42, 3.46 and 4.46. c. In the inactive structure (PDB code 5U09, grey), residue 3.46 which is Thr in WT was mutated to Ala (coloured green) to aid in structural determination. This inactivating T210^3.46^A mutation is close to F155^2.42^ and might influence its conformation.

**Fig S5.**
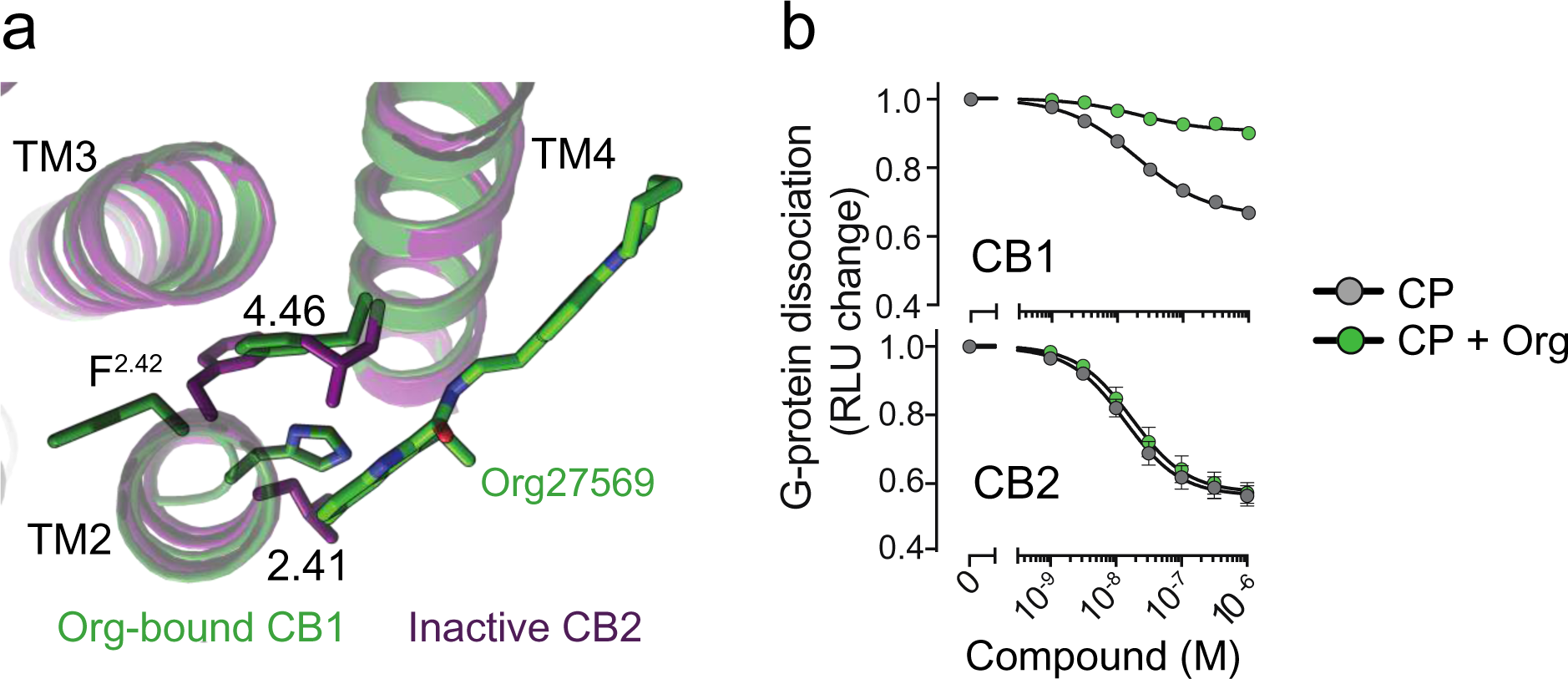
a. Structure of Org-bound CB1 (PDB: 6KQI, green) showing interaction with H154^2.41^. In CB2 (purple) position 2.41 is a Leu, which might prevent Org binding to CB2. b. NanoBiT-G-protein dissociation assay shows unchanged CP response upon Org treatment in CB2.

**Table S1:**
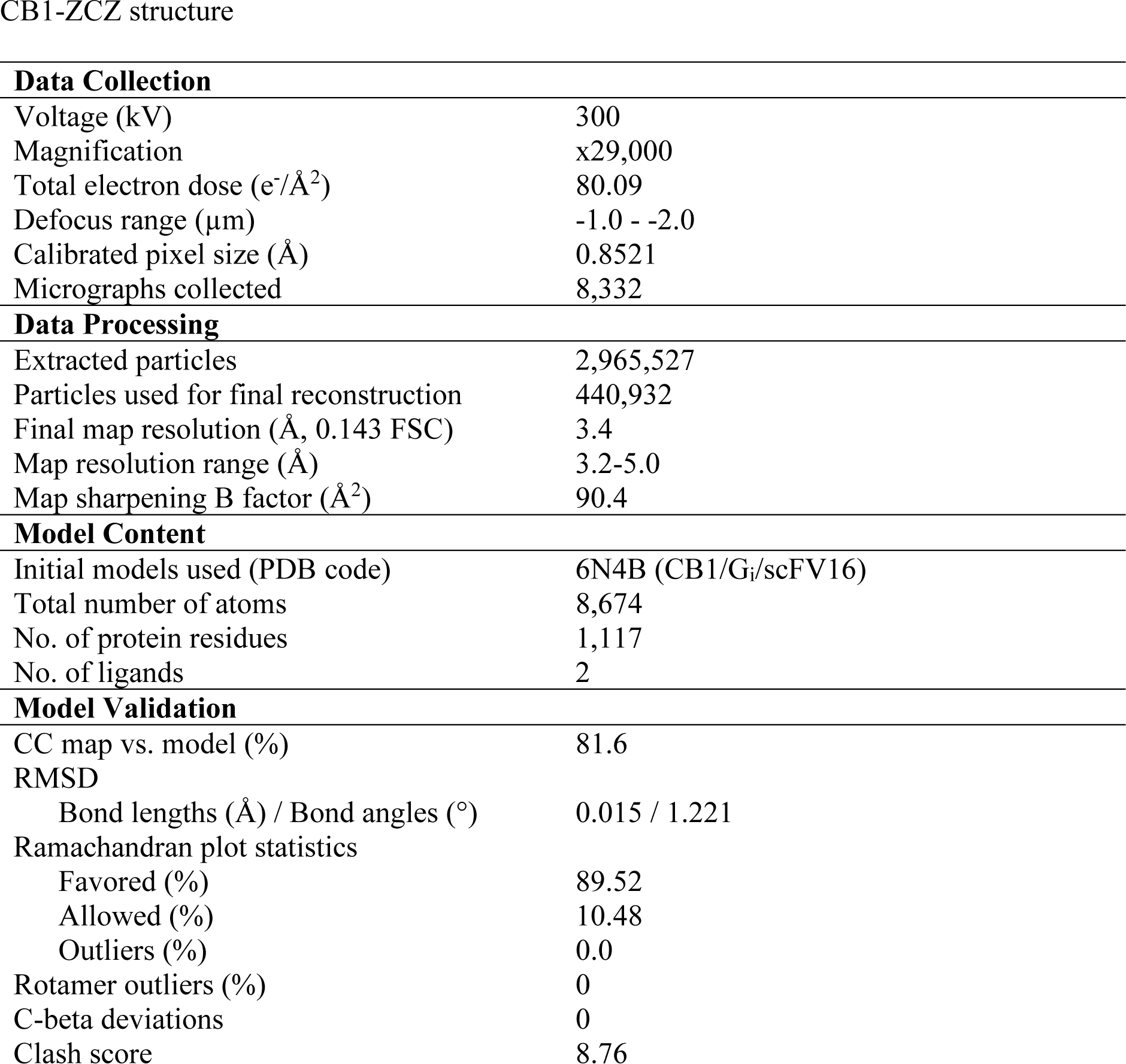
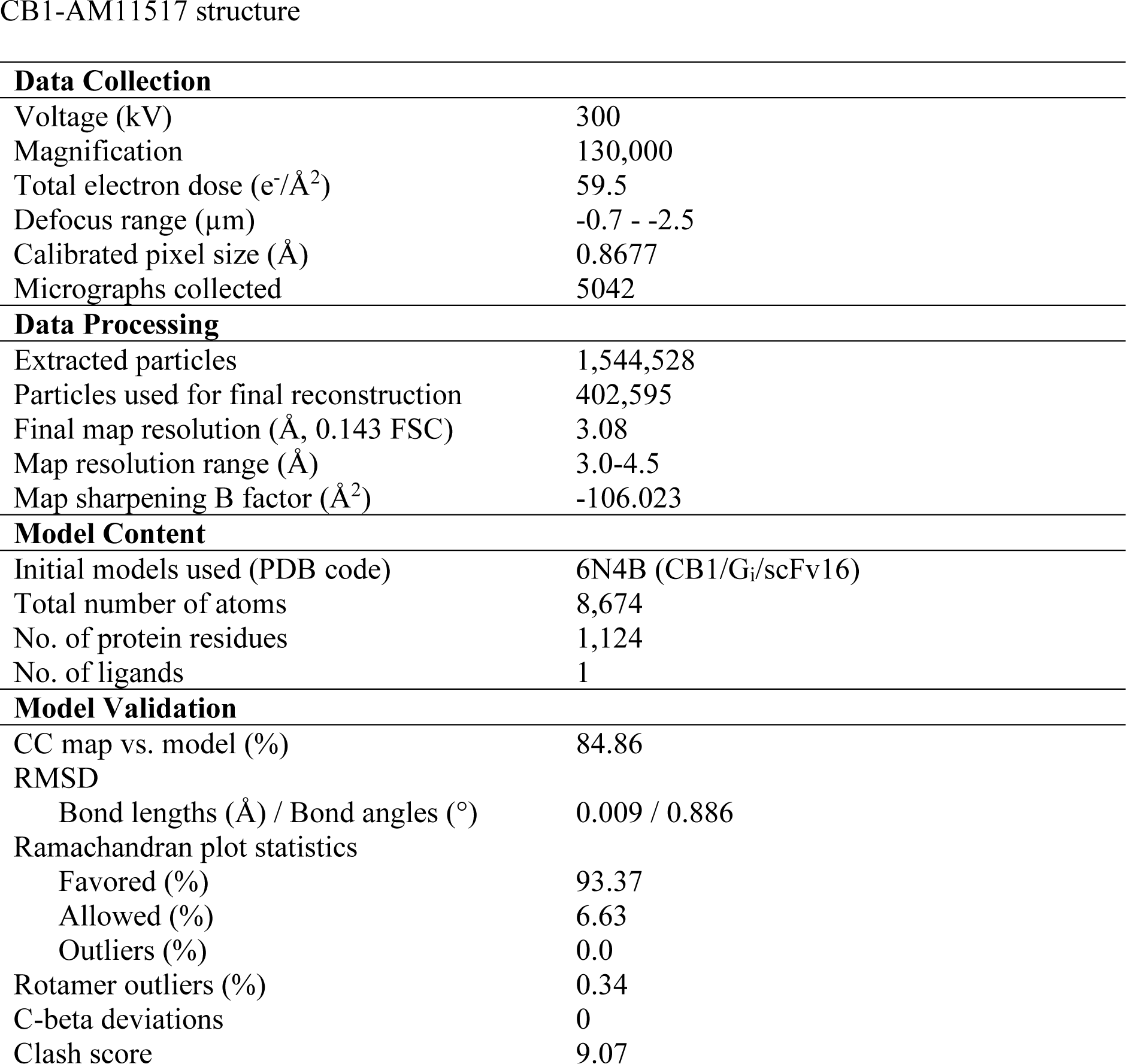
CryoEM data collection, model refinement and validation.

CB1-ZCZ structure
| **Data Collection** |  |
| --- | --- |
| Voltage (kV) | 300 |
| Magnification | x29,000 |
| Total electron dose (e<sup>-</sup>/Å<sup>2</sup>) | 80.09 |
| Defocus range (μm) | -1.0 - -2.0 |
| Calibrated pixel size (Å) | 0.8521 |
| Micrographs collected | 8,332 |
| **Data Processing** |  |
| Extracted particles | 2,965,527 |
| Particles used for final reconstruction | 440,932 |
| Final map resolution (Å, 0.143 FSC) | 3.4 |
| Map resolution range (Å) | 3.2-5.0 |
| Map sharpening B factor (Å<sup>2</sup>) | 90.4 |
| **Model Content** |  |
| Initial models used (PDB code) | 6N4B (CB1/Gi/scFV16) |
| Total number of atoms | 8,674 |
| No. of protein residues | 1,117 |
| No. of ligands | 2 |
| **Model Validation** |  |
| CC map vs. model (%) | 81.6 |
| RMSD |  |
| Bond lengths (Å) / Bond angles (°) | 0.015 / 1.221 |
| Ramachandran plot statistics |  |
| Favored (%) | 89.52 |
| Allowed (%) | 10.48 |
| Outliers (%) | 0.0 |
| Rotamer outliers (%) | 0 |
| C-beta deviations | 0 |
| Clash score | 8.76 |

**Table S1: CryoEM data collection, model refinement and validation** **CB1-AM11517 structure**
| **Data Collection** |  |
| --- | --- |
| Voltage (kV) | 300 |
| Magnification | 130,000 |
| Total electron dose (e<sup>-</sup>/Å<sup>2</sup>) | 59.5 |
| Defocus range (μm) | -0.7 - -2.5 |
| Calibrated pixel size (Å) | 0.8677 |
| Micrographs collected | 5042 |
| **Data Processing** |  |
| Extracted particles | 1,544,528 |
| Particles used for final reconstruction | 402,595 |
| Final map resolution (Å, 0.143 FSC) | 3.08 |
| Map resolution range (Å) | 3.0-4.5 |
| Map sharpening B factor (Å<sup>2</sup>) | -106.023 |
| **Model Content** |  |
| Initial models used (PDB code) | 6N4B (CB1/Gi/scFv16) |
| Total number of atoms | 8,674 |
| No. of protein residues | 1,124 |
| No. of ligands | 1 |
| **Model Validation** |  |
| CC map vs. model (%) | 84.86 |
| RMSD |  |
| Bond lengths (Å) / Bond angles (°) | 0.009 / 0.886 |
| Ramachandran plot statistics |  |
| Favored (%) | 93.37 |
| Allowed (%) | 6.63 |
| Outliers (%) | 0.0 |
| Rotamer outliers (%) | 0.34 |
| C-beta deviations | 0 |
| Clash score | 9.07 |

## Methods

### Purification of CB1

CB1 was expressed and purified as described previously ^6^. Briefly, human full-length CB1 containing a N-terminal FLAG tag and C-terminal histidine tag was expressed in *Spodoptera frugiperda Sf9* insect cells with the baculovirus method (Expression Systems). Receptor was extracted using 1% lauryl maltose neopentyl glycol (L-MNG) and purified by nickel-chelating Sepharose chromatography. The eluant from the Ni column was applied to a M1 anti-FLAG immunoaffinity resin. After washing to progressively decreasing concentration of L-MNG, the receptor was eluted in a buffer consisting of 20 mM HEPES pH 7.5, 150 mM NaCl, 0.05% L- MNG, 0.005% cholesterol hemisuccinate (CHS), FLAG peptide and 5 mM EDTA. Finally, CB1 was purified with size exclusion chromatography, on Superdex 200 10/300 gel filtration column (GE) in 20 mM HEPES pH 7.5, 150 mM NaCl, 0.02% L-MNG, 0.002% CHS. Ligand-free CB1 was concentrated to ∼500 µM and stored in -80 °C.

### Expression and purification of G_i_ heterotrimer

Heterotrimeric G_i_ was expressed and purified as previously described ^23^. Insect cells (*Trichuplusia ni, Hi5*) was co-infected with wild-type human Gα_i_ subunit virus and wild-type human β1γ2 virus. β_1_γ_2_ contains an histidine tag inserted at the amino terminus of the β subunit that is used for further purification. After harvesting cells expressing the heterotrimetric G- protein, they were lysed in hypotonic buffer. Heterotrimeric G_i_β_1_γ_2_ was extracted in a buffer containing 1% sodium cholate and 0.05% n-dodecyl-β-D-maltoside (DDM, Anatrace). Ni-NTA chromatography is performed and the detergent was exchanged from cholate/DDM to DDM on column. After elution, the protein was dialyzed overnight dialyzed overnight in 20 mM HEPES, pH 7.5, 100 mM sodium chloride, 0.1% DDM, 1 mM magnesium chloride, 100uM TCEP and 10 μM GDP together with Hugman rhinovirus 3C protease (3C protease) to cleave off the amino- terminal 6xHis tag. 3C protease was removed by Ni-chelated sepharose and the heterotrimetric G-protein was further purified with MonoQ 10/100 GL column (GE Healthcare). Protein was bound to the column and washed in buffer A (20 mM HEPES, pH 7.5, 50 mM sodium chloride, 1 mM magnesium chloride, 0.05% DDM, 100 μM TCEP, and 10 μM GDP). The protein was eluted with a linear gradient of 0–50% buffer B (buffer A with 1 M NaCl). The collected G protein was dialyzed into 20 mM HEPES, pH 7.5, 100 mM sodium chloride, 1 mM magnesium chloride, 0.02% DDM, 100 μM TCEP, and 10 μM GDP. Protein was concentrated to 250 µM and flash frozen until further use.

### Purification of scFv16

scFv16 was purified with a hexahistidine-tag in the secreted form from *Trichuplusia ni Hi5* insect cells using the baculoviral method. The supernatant from baculoviral infected cells was pH balanced and quenched with chelating agents and loaded onto Ni resin. After washing with 20 mM HEPES pH 7.5, 500 mM NaCl, and 20 mM imidazole, protein was eluted with 250 mM imidazole. Following dialysis with 3C protease into a buffer consisting of 20mM HEPES pH 7.5 and 100 mM NaCl, scFv16 was further purified by reloading over Ni a column. The collected flow-through was applied onto a Superdex 200 16/60 column and the peak fraction was collected, concentrated and flash frozen.

### CB1-G_i_ complex formation and purification

CB1 in L-MNG was incubated with AMG315 and ZCZ (or AM11517) for ∼ 1 hour at room temperature. Simultaneously, G_i_1 heterotrimer in DDM was incubated with 1% L-MNG at 4 °C. The AMG315- and ZCZ (or AM11517)-bound CB1 was incubated with a 1.25 molar excess of detergent exchanged G_i_ heterotrimer at room temperature for ∼ 3 hour. To stabilize a nucleotide-free complex, apyrase was added and incubated for 1.5 hour at 4 °C. The complex was diluted 4-fold with 20 mM HEPES pH 7.5, 100 mM NaCl, 0.8% L-MNG/0.08% CHS, 0.27% GDN/0.027% CHS, 1 mM MgCl2, 10 µM AMG315, 20 µM ZCZ (or AM11517) and 2 mM CaCl2 and purified by M1 anti-FLAG affinity chromatography. After washing to remove excess G protein and reduce detergents, the complex was eluted in 20mM HEPES pH 7.5, 100mM NaCl, 0.01% L-MNG/0.001% CHS, 0.0033% GDN/0.00033% CHS, 10 µM AMG315, 10 µM ZCZ (or AM11517), 5 mM EDTA, and FLAG peptide. The complex was supplemented with 100 µM TCEP and incubated with 2 molar excess of scFv16 overnight at 4 °C. Size exclusion chromatography (Superdex 200 10/300 Increase) was used to further purify the CB1-G_i_-scFv16 complex. The complex in 20mM HEPES pH 7.5, 100mM NaCl, 10 µM AMG315, 10 µM ZCZ (or AM11517), 0.00075% L-MNG/0.000075% CHS and 0.00025% GDN/0.000025% CHS was concentrated to ∼15 mg/mL for electron microscopy studies.

### Cryo-EM data acquisition

#### AMG315/ZCZ/CB1/Gi Complex Structure

For grid preparation, 3 μL of purified CB1-G_i_ complex at 15 mg/ml was applied on glow-discharged holey carbon gold grids (Quantifoil R1.2/1.3, 200 mesh). The grids were blotted using a Vitrobot Mark IV (FEI) with 4 s blotting time and blot force 3 at 100% humidity and plunge-frozen in liquid ethane. A total of 8332 movies were recorded on a Titan Krios electron microscope (Thermo Fisher Scientific- FEI) operating at 300 kV at a calibrated magnification of x29,000 and corresponding to a pixel size of 0.8521 Å. Micrographs were recorded using a K3 Summit direct electron camera (Gatan Inc.) with a dose rate of 1.405 electrons/Å^2^/s. The total exposure time was 3.895 s with an accumulated dose of ∼80.09 electrons per Å^2^ and a total of 57 frames per micrograph. Automatic data acquisition was done using *SerialEM*.

### AM11517-bound structure

For grid preparation, 3.5 μL of purified CB1-G_i_ complex at 12 mg/ml was applied on glow-discharged holey carbon gold grids (Quantifoil R1.2/1.3, 200 mesh) at room temperature (for AM11517). The grids were blotted using a Vitrobot Mark IV (FEI) with 3 s blotting time at 100% humidity and plunge-frozen in liquid ethane. A total of 4,837 movies were recorded on a Titan Krios electron microscope (Thermo Fisher Scientific- FEI) operating at 300 kV at a calibrated magnification of x29,000 and corresponding to a magnified pixel size of 0.86 Å. Micrographs were recorded using a K3 Summit direct electron camera (Gatan Inc.) with a dose rate of ∼5.0 electrons/Å^2^/s and defocus values ranging from -0.7μm to -2.0 μm. The total exposure time was 10.0 s and intermediate frames were recorded in 0.2 s intervals resulting in an accumulated dose of ∼51.65 electrons per Å^2^ and a total of 50 frames per micrograph. Automatic data acquisition was done using *SerialEM*.

### Image processing and 3D reconstructions

#### AMG315/ZCZ/CB1/Gi Complex Structure

For the dataset of the AMG315/ZCZ/CB1/Gi complex, micrographs were imported into RELION 3.1 and beam-induced motion correction was performed with *MotionCor2* followed by CTF parameter fitting with *CTFFIND4*. Extracted particles were sorted with iterative rounds of 2D classification followed by iterative rounds of 3D classification to arrive at a final curated stack of 440,932 particles. These particles were then subjected to Bayesian polishing ^24^ and final reconstruction in RELION 3.1 (**Fig. S1b**).

#### AM11517-bound structure

Micrographs were subjected to beam-induced motion correction using *MotionCor2* ^25^. CTF parameters for each micrograph were determined by *CTFFIND4* ^26^. An initial set of 1,544,528 particle projections were extracted using semi-automated procedures and subjected to reference-free two-dimensional classification in *RELION 2.1.0* ^27^. From this step, 562,312 particle projections were selected for further processing. The map of CB1 receptor low passed filtered to 60 Å was used as an initial reference model for maximum-likelihood-based three- dimensional classifications. Conformationally homogeneous groups accounting for 177,787 particles, forming class averages with well resolved features for all subunits, were subjected to 3D masked refinement in *Frealign* (*CisTEM* ^28^) followed by map sharpening applying temperature-factors of -90 Å2 and -60 Å2 for the low- and high- resolution ends of the amplitude spectrum, respectively. The final map has an indicated global nominal resolution of 3.1 Å (**Fig. S1d**). Reported resolution is based on the gold-standard Fourier shell correlation (FSC) using the 0.143 criterion and is in agreement with both Relion 2.1.0 and *M-triage* as implemented in *Phenix* ^29^. Local resolution was determined using *B-soft* ^30^ with half map reconstructions as input maps (**Fig. S1d**).

#### Model building and refinement

The initial template of CB1 was the FUB-bound CB1-G_i_ structure (PDB 6n4b). Agonist coordinates and geometry restrains were generated using *phenix.elbow.* Models were docked into the EM density map using *UCSF*. *Coot* was used for iterative model building and the final model was subjected to global refinement and minimization in real space using *phenix.real_space_refine* in *Phenix*. Model geometry was evaluated using *Molprobity*. FSC curves were calculated between the resulting model and the half map used for refinement as well as between the resulting model and the other half map for cross-validation (**Fig. S1b, d**). The final refinement parameters are provided in **Table S1**.

#### GTP turnover assay

Analysis of GTP turnover was performed by using a modified protocol of the GTPase-Glo^TM^ assay (Promega) described previously ^31^. Unliganded or liganded-CB1 (1 uM) and G_i_ (1 uM) was mixed together in 20 mM HEPES, pH 7.5, 50 mM NaCl, 0.01% L-MNG, 100 μM TCEP, 10 μM GDP and 5 μM GTP. GTPase-Glo-reagent was added to the sample after incubation for 60 minutes (agonist assays) or 30 minutes (for PAM assays). Luminescence was measured after the addition of detection reagent and incubation for 10 min at room temperature using a *SpectraMax Paradigm* plate reader.

### MD simulations

#### System Setup for MD Simulation

We performed simulations of CB1R bound to the endocannabinoid Anandamide, to the synthetic cannabinoid FUB, and to AMG315, an analogue of Anandamide. We initiated the simulations from the AMG315-bound structure that was solved in the presence of ZCZ presented in this paper. For all simulations, we removed the single chain variable fragment (scFv) and the G protein from the structure. For the FUB-bound and Anandamide-bound simulations, we replaced the AMG315 molecule with FUB or Anandamide in silico in Maestro (Schrödinger). For each of these three simulation conditions, we performed six independent simulations in which initial atom velocities were assigned randomly and independently.

Neutral acetyl and methylamide groups were added to cap the N- and C-termini, respectively, of protein chains. Extracellular loop 2 (ECL2) loop of the receptor was modeled using the Maestro (Schrödinger) “crosslinking” tool with a fragment from the previously published structure of CB1 bound to agonist AM11542 (PDB ID: 5XRA) ^15^. Titratable residues were kept in their dominant protonation states at pH 7, except for D2.50 (D163) and D3.49 (D213), which were protonated (neutral) in all simulations, as studies indicate that these conserved residues are protonated in active-state GPCRs^32, 33^. Histidine residues were modeled as neutral, with a hydrogen bound to either the delta or epsilon nitrogen depending on which tautomeric state optimized the local hydrogen-bonding network. Dowser was used to add water molecules to protein cavities, and the protein structures were aligned on transmembrane (TM) helices of the FUB-bound active CB1 crystal structure (PDB ID: 6N4B) ^6^ in the Orientation of Proteins in Membranes (OPM) database ^34^. The aligned structures were inserted into a pre-equilibrated palmitoyl-oleoyl-phosphatidylcholine (POPC) membrane bilayer using Dabble ^35^. Sodium and chloride ions were added to neutralize each system at a concentration of 150 mM. Systems comprised 56,000 atoms, including ∼140 lipid molecules and ∼11,000 water molecules. Approximate system dimensions were 80 Å x 90 Å x 85 Å.

#### Simulation Protocols

Simulations were run using the AMBER18 software ^36^ under periodic boundary conditions with the Compute Unified Device Architecture (CUDA) version of Particle-Mesh Ewald Molecular Dynamics (PMEMD) on graphics processing units (GPUs) ^37^. The systems were first heated over 12.5 ps from 0 K to 100 K in the NVT ensemble using a Langevin thermostat with harmonic restraints of 10.0 kcal·mol-1·Å^-2^ on the non-hydrogen atoms of the lipids, protein, and ligand. Initial velocities were sampled from a Boltzmann distribution. The systems were then heated to 310 K over 125 ps in the NPT ensemble. Equilibration was performed at 310 K and 1 bar in the NPT ensemble, with harmonic restraints on the protein and ligand non-hydrogen atoms tapered off by 1.0 kcal·mol-1 ·Å^-2^ starting at 5.0 kcal·mol-1 ·Å^-2^ in a stepwise manner every 2 ns for 10 ns, and finally by 0.1 kcal·mol-1 ·Å^-2^ every 2 ns for an additional 18 ns. All restraints were completely removed during production simulation. Production simulations were performed at 310 K and 1 bar in the NPT ensemble using the Langevin thermostat and Monte Carlo barostat. The simulations were performed using a timestep of 4.0 fs while employing hydrogen mass repartitioning. Bond lengths were constrained using SHAKE. Non-bonded interactions were cut off at 9.0 Å, and long-range electrostatic interactions were calculated using the particle-mesh Ewald (PME) method with an Ewald coefficient (β) of approximately 0.31 Å and B-spline interpolation of order 4. The PME grid size was chosen such that the width of a grid cell was approximately 1 Å. We employed the CHARMM36m force field for protein molecules, the CHARMM36 parameter set for lipid molecules and salt ions, and the associated CHARMM TIP3P model for water ^38, 39^. Ligand parameters were obtained using the CGenFF webserver ^40, 41^.

For each ligand, we performed 6 independent 2-µs simulations at 310 K. All simulations were performed on the Sherlock computing cluster at Stanford University.

#### Simulation Analysis Protocols

The AmberTools17 CPPTRAJ package was used to reimage trajectories at 1 ns per frame, while Visual Molecular Dynamics (VMD) ^42^was used for visualization and analysis. For all reported analyses, we discarded the first 0.5 µs of each simulation to achieve better equilibration.

For Figure 3, we determined the fraction of time W6.48 (W356) spent in an active-like conformation by setting a threshold value of 6.7 Å for the distance between the beta carbon of W6.48 and the alpha carbon of C7.42 on TM7. Frames with a distance greater than the threshold were classified as active-like. To determine whether differences between simulations performed with different ligands were statistically significant, we performed two-sided t-tests of unequal variance (Welch’s t-test) on the frequency of this distance being above the threshold value, with each simulation as an independent sample.

For Figure 4, we used GetContacts (https://getcontacts.github.io/) to determine frequency of polar interactions between each ligand and H2.65 (H178) in simulation. Specific polar contacts considered were direct hydrogen bonds or hydrogen bonds mediated by one water molecule. To determine whether differences between simulation conditions performed with different ligands were statistically significant, we performed two-sided t-tests of unequal variance (Welch’s t-test) on the frequency of polar interactions using each simulation as an independent sample.

#### NanoBiT-G-protein dissociation assay

CB1-induced G-protein dissociation was measured by the NanoBiT-G-protein dissociation assay, in which the interaction between the Gα subunit and the Gγ subunit was monitored by the NanoBiT-based enzyme complementation system (Promega). Specifically, the NanoBiT- G_i_1 protein consisting of the Gαi1 subunit fused with a large fragment (LgBiT) at the α-helical domain and the N-terminally small fragment (SmBiT)-fused Gγ2 subunit was expressed, along with an untagged Gβ1 subunit and a test GPCR construct. CB1 construct with the N-terminal hemagglutinin signal sequence and the FLAG epitope tag with a flexible linker (MKTIIALSYIFCLVFADYKDDDDKGGSGGGGSGGSSSGGG) was inserted into the pCAGGS expression vector. HEK293A cells (Thermo Fisher Scientific) were seeded in a 10- cm culture dish at a concentration of 2 x 10^5^ cells ml^-1^ (10 ml per dish in DMEM (Nissui) supplemented with 10% fetal bovine serum (Gibco), glutamine, penicillin and streptomycin), one day before transfection. The transfection solution was prepared by combining 25 µl (per dish hereafter) of polyethylenimine (PEI) Max solution (1 mg ml^-1^; Polysciences), 1 ml of Opti-MEM (Thermo Fisher Scientific) and a plasmid mixture consisting of 1 µg test GPCR construct, 500 ng LgBiT-containing Gαi1 subunit, 2.5 µg Gβ1 subunit and 2.5 µg SmBiT- fused Gγ2 subunit with the C68S mutant. After an incubation for one day, the transfected cells were harvested with 0.5 mM EDTA-containing Dulbecco’s PBS, centrifuged, and suspended in 9 ml of HBSS containing 0.01% bovine serum albumin (BSA; fatty acid–free grade; SERVA) and 5 mM HEPES (pH 7.4) (assay buffer). The cell suspension was dispensed in a white 96-well plate at a volume of 70 µl per well and loaded with 20 µl of 50 µM coelenterazine (Carbosynth) diluted in the assay buffer. After a 2 h incubation at room temperature, the plate was measured for baseline luminescence (SpectraMax L, Molecular Devices) and a test allosteric ligand (10 µl) was manually added. The plate was immediately read at room temperature for the following 10 min as the kinetics mode, at measurement intervals of 20 sec. Thereafter, a test orthosteric ligand (20 µl) was added and the plate was read for another 10 min. The luminescence counts over 3-5 min after ligand addition were averaged and normalized to the initial count. The fold-change values were further normalized to that of vehicle-treated samples, and used to plot the G-protein dissociation response. Using the Prism 8 software (GraphPad Prism), the G-protein dissociation signals were fitted to a four-parameter sigmoidal concentration-response curve, from which pEC50 values (negative logarithmic values of EC50 values) and E_max_ values were used to calculate mean and SEM.

## Data availability

The cryo-EM density maps has been deposited in the Electron Microscopy Data Bank (EMDB) under accession code EMD-XXXX and EMD-XXXX. Model coordinates have been deposited in the Protein Data Bank (PDB) under accession number XXXX and XXXX.

## Author Contributions

K.K. prepared the CB1-G_i_ complex, collected cryo-EM data, processed data and obtained the cryo-EM map (AM11517-bound CB1 complex) with help from H.W. Modelled both the structures. Performed the GTP-turnover assays.

M.J.R collected cryo-EM data, processed data and obtained the cryo-EM map for ZCZ-bound CB1 complex under supervision of G.S. Performed docking for the AMG315 enantiomer. E.T., C.-M.S, and A.S.P. performed and analyzed molecular dynamics simulations under supervision of R.O.D.

A.S. performed the NanoBiT assay.

L.J., S.P.N, M.G. and K.V. designed and synthesized ligands supervised by A.M.

K.K. and B.K.K. wrote the manuscript with input from all the authors.

## Acknowledgements

Kaavya Krishna Kumar was supported by the ADA Fellowship. C.-M.S. is supported by Human Frontier Science Program Long-Term Fellowship LT000916/2018-L. R.O.D acknowledges National Institutes of Health for grant R01GM127359. We would like to thank National Institute of Drug Abuse (NIDA). B.K.K. is a Chan Zuckerberg Biohub investigator.

